# Semi-automated retrieval of chemical and phylogenetic information from natural products literature

**DOI:** 10.1101/2023.06.28.546864

**Authors:** Ana Carolina Lunardello Coelho, Ricardo R. da Silva

## Abstract

Natural products (NPs) are metabolites of great importance due to their fundamental biological role in performing specialized activities, ranging from basic cellular functions to complex ecological interactions. These metabolites have contributed to innovating fields such as agriculture and medicine due to their optimized biological activities, a consequence of evolution. A key factor in ensuring that isolated NPs are novel is to search scientific literature and compare pre-existing chemical entities with the new isolate. Unfortunately, articles are typically not machine-readable, a problem that hinders efficient searching and increases the chances of unintended rediscovery. In addition, the time required to add new compound discoveries to compound databases hinders computational studies on cell metabolism and Quantitative Structure-Activity Relationships (QSAR). Here, we present a modularized tool that uses text mining techniques to retrieve chemical entities and taxonomic mentions present in scientific literature, called NPMINE (Natural Products MINIng). We were able to analyze 55,382 scientific articles from some of the most important applied chemistry journals from Brazil and the world, consistently recovering the expected taxonomic and structural information. This processing resulted in 120,970 unique InChI Keys potentially associated with 21,526 unique species mentioned. Using the PubChem BioAssay database we show how QSAR models can be used to mine active leads. The results indicate that NPMINE not only facilitates natural products cataloging, but also assists in biological source assignment and structure-activity relationships, a time-consuming task, typically performed in low throughput.

## Introduction

Natural products (NPs) are specialized metabolites with biotechnological applications present in various fields such as medicine, agriculture and cosmetics [1–4]. These metabolites have gained the interest of the scientific community due to their bioactivity and complex chemical structures [5, 6]. Over the years, many novel NPs have been discovered by isolating compounds of interest. These discoveries lead to technological innovation and socio-economic growth.

A challenge many scientists face when researching NPs is determinating if their isolate isn’t a rediscovery, a process also referred as dereplication. Typically, dereplication is performed by searching scientific literature for potential matches with the isolate being studied. New NPs are generally described by a non-standard name, that usually refers to the source of isolation. These names are not informative about the structure of the compound and may be redundant, hampering cataloging and future application. Image representations of chemical structure, which are defined as unstructured data types and are not machine-readable, may also be present in scientific articles [7]. Thus, the results of searching for potential matches tends to be non-exhaustive due to the multiple ways in which a chemical compound can be named [8], and by its presence as images embedded in scientific articles. The DECIMER (Deep lEarning for Chemical IMagE Recognition) Segmentation [9], OC processor [10], OSRA (Optical Structure Recognition Application) [11] and ChemDataExtractor [7] are all examples of tools designed to assist in this endeavor by extracting unstructured or semi-structured chemical entities.

Web applications and databases focused on NPs have been developed [12] to catalog, search and study NPs. Some are compilations of previously described natural product databases [13, 14], others specialize in NPs coming from specific sources [15, 16], focus on providing the compound and its biological origin [17, 18], or specialize in NPs present in the biodiversity of a specific region [19, 20]. One of the most popular databases, the Dictionary of Natural Products (DNP), is only commercially available, which limits development in the field. Most recently, the COCONUT (COlleCtion of Open Natural ProdUcTs) [13] and LOTUS (naturaL prOducTs occUrrence databaSe) databases [21] were created as an attempt to solve these issues by developing an open source, cross kingdom, comprehensive machine-readable resource that links NPs to the organisms which produce them and also allows for user contributions. Additionally, NUBBEDB (Nuclei of Bioassays, Biosynthesis and Ecophysiology of Natural Products Database) [19] and NPAtlas (Natural Products Atlas) [15], are examples of databases that were created, at least in part, through manual curation, which is a very time consuming task that could be facilitated by a tool streamlining cataloging.

Therefore, NPMINE (Natural Product MINing), a Python library, was developed as a tool for the scientific community to easily find NPs present in scientific articles. Third party tools such as OSRA are used to extract chemical information from scientific articles that are in text and image formats. The semi-automated nature of NPMINE allows scientists to build and curate their own in-house databases more efficiently, thus aiding in the process of dereplication and cataloging. In addition to extracting chemical compounds, NPMINE also retrieves taxonomic information present in the text and can help to find metabolite-biological source pairings of interest.

## Results and Discussion

### NPMINE workflow

NPMINE is a Python library that processes the outputs of third party tools, as described in detail in later sections of this paper, and its workflow can be visualized in Figure 1. The third party tools and NPMINE are provided as a docker image for ease of use. We also provide a comprehensive repository (https://github.com/computational-chemical-biology/npmine), with source code for NPMINE, it’s docker image, a snakemake script [22] for easy management of the complete workflow and a set of example Jupyter notebooks, aimed at showcasing the package utilities as well as making this work fully reproducible. The workflow starts with the extraction of the article URL link from the journal databases found online, either by using web scraping techniques or by using the API (Application Program Interface) access provided by the journal. Articles stored locally on the user’s system can also be processed by NPMINE by using the notebook examples available in the repository.

**Figure 1.**
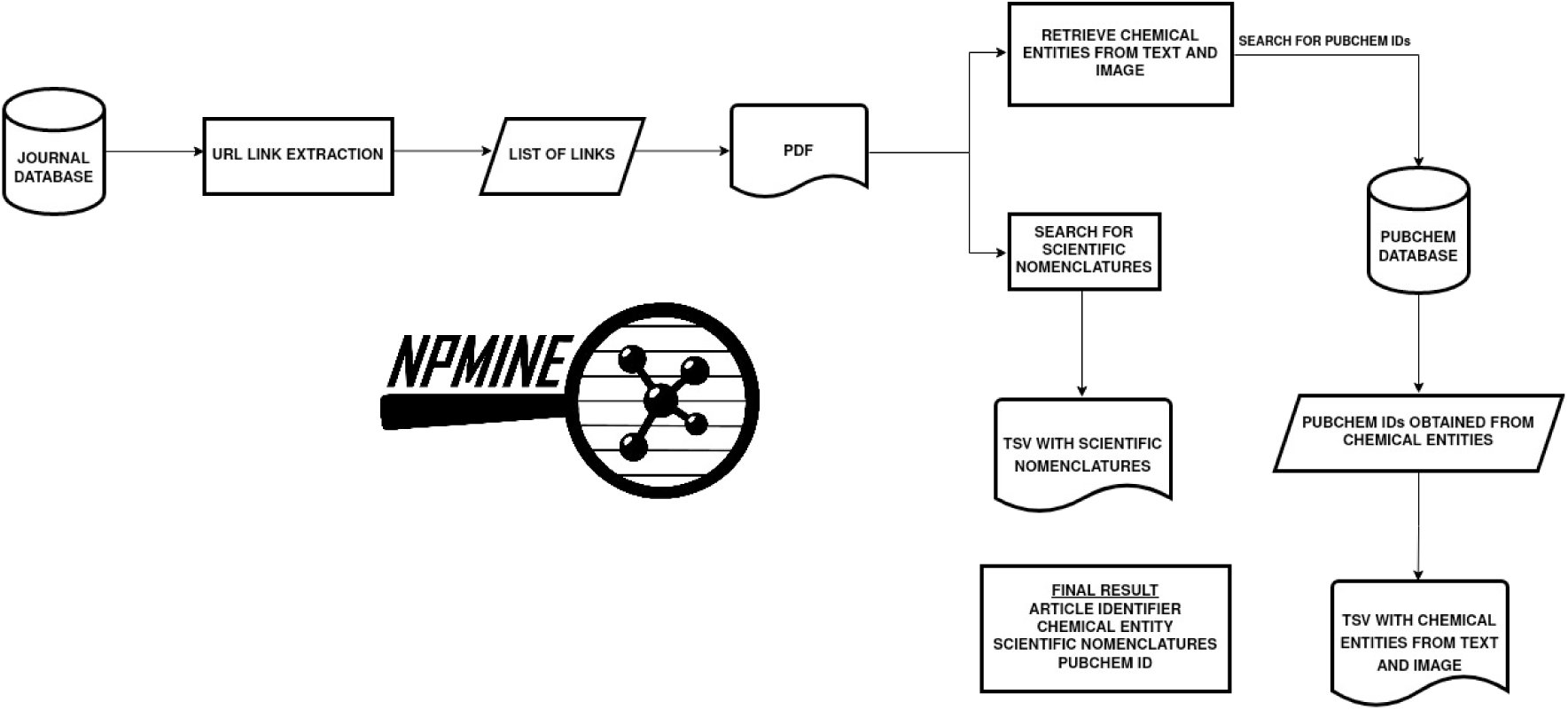
NPMINE’s Workflow. The link of each article was extracted from the journal databases. This list was used to download the articles. Chemical entities present in the PDF were retrieved. The InChI obtained were then converted to InChIKeys and used to query PubChem. The scientific names present were also retrieved. The final result is summarized in two TSV outputs.

Given a collection of PDF documents, chemical entities can be retrieved from the text using Chemextractor, which uses OSCAR4 (Open Source Chemistry Analysis Routines) [23], or from images, using OSRA (Optical Structure Recognition Application) [11]. Once the chemical information is retrieved, if it is in the form of InChI or SMILES, it can then be converted to InChI Keys and used to query PubChem and retrieve database IDs, if available. Scientific names, retrieved by Gnfinder (Global Names Finder) [24], are also extracted from the text. The final result is a summary of the chemical and taxonomic information retrieved, associated with a unique identifier for each journal article.

### Benchmarking NPMINE

To quantify NPMINE’s ability to recover chemical and taxonomic information, we performed a benchmarking experiment (Fig. 2-A). The experiment consisted of simulating PDF documents with chemical and taxonomic information in the form of text and chemical information in the form of embedded images. A LATEX [25] template was used to store the information randomly selected from the Super Natural (SUP) [14], a NPs database, and scientific names from the NCBI (National Center for Biotechnology Information) taxonomy database [26]. These randomly selected information were mixed with simulated text and images, formatted as sections of a typical scientific article, using an American Chemical Society template. An example can be found included in Additional file 1: Fig. S1. All randomly selected information was stored for comparison with the information retrieved by NPMINE. The script *generate_pdf* is part of the NPMINE library, allowing the reproduction of this workflow. Using this approach we simulated a thousand documents, and applied NPMINE to automatically recover the information. The most challenging task performed by NPMINE is the recovery of chemical structures embedded in images. The OSRA package is amongst one of the best performing open source tools available to this task [27], however, many of the structures recovered are not identical to the ones used to generate the images (Additional file 2: Fig. S2). In our simulation we observed that 46% of the recovered structures had a similarity score equal or higher than 0.5 (Fig. 2-B). Although it may indicate a limited ability of OSRA to retrieve compounds, the recovered structures sometimes differ in small structural features, which can be easily fixed by using free software for chemical structure drawing, speeding the process to discover and catalog compounds, otherwise performed entirely manually. An easy to access interface linking the original document from which the structure was recovered to PubChem Sketcher [28]. We have created an output that easily connects the recovered structure to the document as well as to a structure drawing software, allowing corrections if necessary [See Additional file 5 : Fig. S4].

**Figure 2.**
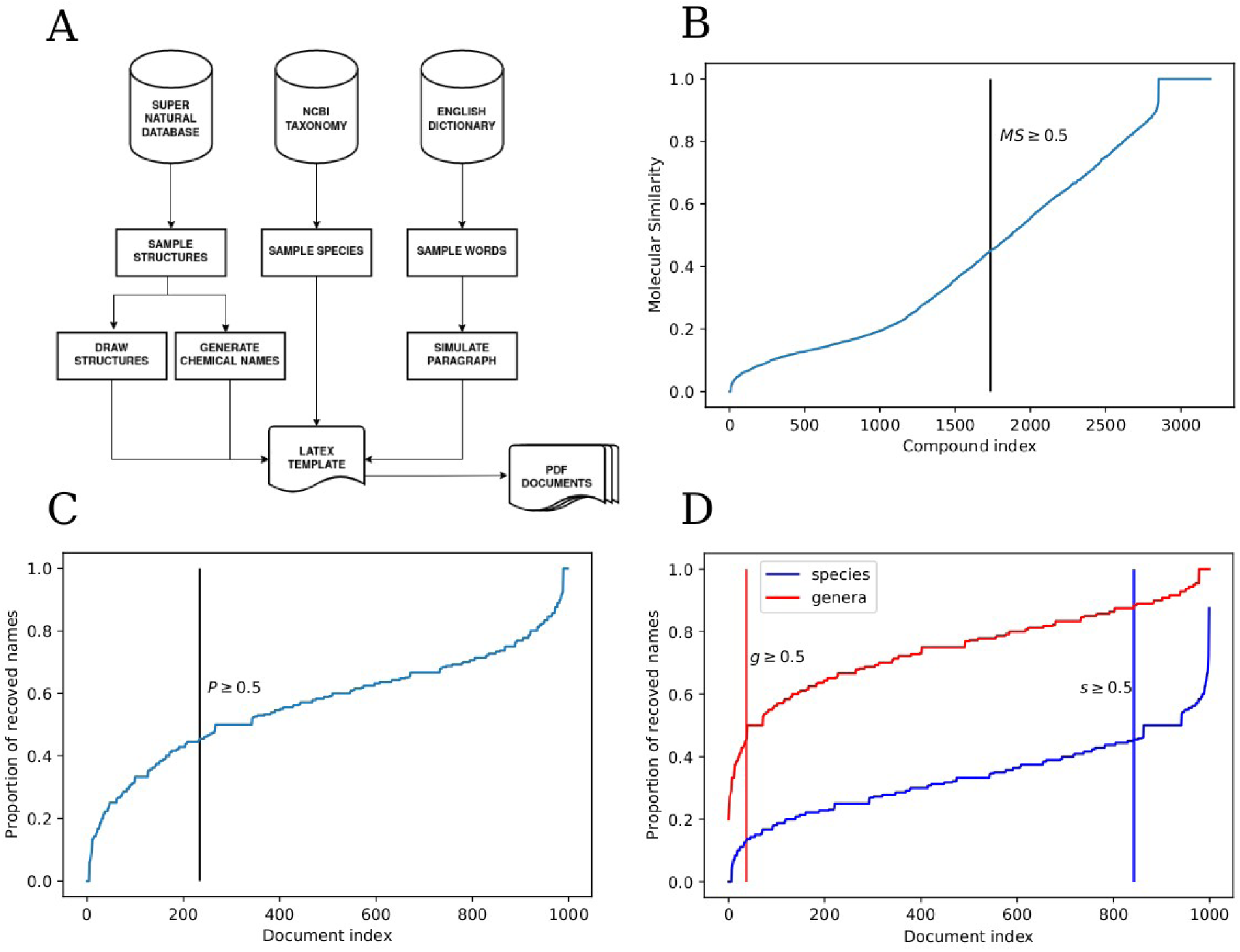
Benchmarking experiment for NPMINE’s Workflow. A - Workflow describing each step of the benchmarking experiment. B - Molecular similarity between the compounds included in the experiments, with the version recovered by NPMINE (through OSRA). C - Proportion of chemical names recovered (through OSCAR4) within each document. D - Proportion of scientific names recovered (through Gnfinder) within each document, at genera and species level.

The ambiguity of common chemical names or identifiers is well known in chemical literature [8, 29]. The common name is usually preferred over the IUPAC compound name in NPs literature [30]. OSCAR4 uses a custom Tokeniser to create a list of Token objects, which is a string of characters that correspond to words, to subsequently pass through a ChemNameDictRegistry to create a list of recognized chemical entities, recovering common names, as well as IUPAC names. We used the CACTUS API [31] to translate the SMILES into chemical names and embed them in our simulated documents. NPMINE was able to recover 50% or more of the correct IUPAC names from 76.6% of the documents evaluated (Fig 2-C). For scientific names, Gnfinder was able to recover 50% or more of the correct taxonomic hierarchy down to the species level for only 16.3% of the documents (Fig. 2-D). However, when considered at the genus level this number raised for 96.3% of the documents (Fig. 2-D).

Although incomplete (as shown by the benchmarking results), the information recovery has the potential to aid NPs cataloging. When a taxonomic genus or a partial structure is recovered and associated with a document identifier and link, it speeds the discovery and recording process, allowing decreased manual curation. Furthermore, the creation of rational benchmarking workflows allows the possibility of identifying opportunities of improvement by providing a structured approach to compare performance and, consequently, improving decision-making.

### Applying NPMINE to Natural Products related literature

To further investigate the potential of NPMINE to recover information about NPs, we analyzed 55,382 scientific articles, from open access (*Química Nova*) and five other journals through institutional paid subscription access (Table 1). Most journals have a main web page with links directing to all the editions, issues and articles. These pages allowed us to manually estimate the number of documents contained in each issue and confirm by comparison that our web scraping techniques recovered the links for all articles. Most of the links correspond to article documents, since our scraping technique described above is generic, the HTML DOM (Document Object Model) can also have other links directing to multiple sources, including unstructured pages, which would be recorded by this approach. For example, *Química Nova* has links pointing to documents of different formats (*.doc, .mp4*, for supplementary material, for example). Therefore, document extensions other than *.pdf* were removed before NPMINE processing.

**Table 1.** Chemical and taxonomic information retrieved from each journal applying NPMINE.

| Journal | Number of Articles | Number of InChI Keys | Number of PubChem IDs | Number of Families | Number of Species |
| --- | --- | --- | --- | --- | --- |
| Journal of Natural Products | 14058 | 53005 | 31562 | 841 | 13264 |
| Metabolomics | 1409 | 6283 | 5449 | 116 | 1562 |
| Phytochemistry Reviews | 774 | 14320 | 9598 | 548 | 8595 |
| Química Nova | 6153 | 17945 | 11187 | 294 | 3540 |
| Journal of Chromatography A | 22786 | 36044 | 29055 | 159 | 2900 |
| Journal of Chromatography B | 10202 | 25927 | 18364 | 216 | 2961 |
| Total | 55382 | 120970 | 73428 | 1153 | 21526 |

From the processing of NPMINE we obtained 120,970 unique structures (unique InChI Keys) (Table 1). The processed output retains the source of structure as OSRA, if derived from image, and OSCAR, if derived from text name and can be found in Additional file 3: Table S1. The number of structures recovered and the proportion of those present in PubChem, one of the largest public structure databases, is related to the number of documents as well as the publication scope of each journal. As a journal dedicated to the study of NPs the Journal of Natural Products (JNP), has the highest proportion of recovered InChI Keys (43.81%) and PubChem IDs (42.98%). The JNP has also the lowest proportion of InChI Keys present in PubChem (59.5%), showing the highest potential for discovery of structures not yet fully explored. The Metabolomics journal has the lowest proportion of InChI Keys (5.19%) and the highest proportion of structures deposited in PubChem 86.72%. This result is expected, as Metabolomics journal publishes work mostly interested on the quantitative changes of metabolites, and annotation of known metabolites, in opposition to structural elucidation.

Phytochemistry Reviews has the highest number of InChI Keys by article (18.50%), coherent with a journal publishing literature reviews. The open access Brazilian journal *Química Nova* represents a low proportion of InChI Keys (14.83%), however, it has the second lowest proportion of structures in PubChem (62.34%). This also represents a great potential for cataloging, given the large Brazilian biodiversity and the fact that the articles are published in Portuguese, making them less studied by a broader scientific community. The Journals of Chromatography A and B, which occupy a publication niche for methodological development, have together the largest number of articles (32,988), however, these journals present the lowest potential for discovery of structures not present in public databases, 80.60% and 70.82% of InChI Keys present PubChem, respectively.

### Searching for bioactive leads

In order to investigate the bioactivity potential of compounds recovered by NPMINE, we searched the PubChem BioAssay database for previously reported bioactivity. Five Assays containing more than a thousand compounds recovered by NPMINE were selected to train QSAR models with molecular descriptors as features, using the random forest classifier (Fig. 3-A) [32]. The selected Assays were identified by the Assay ID (AID) and were further filtered by ’Bioactivity Outcome’ to keep only compounds either ‘Active’ or ‘Inactive’ (removing ‘Inconclusive’ and ‘Unspecified’). The trained models had AUCs (Area Under the Curve) ranging from 0.81 to 0.87 and can be found in Additional file 4: Fig. S3. In parallel to the model training, a network of structural similarities between the 120,970 compounds was built, using Morgan Fingerprint to calculate the Tanimoto Similarity. Edges with a similarity below 0.6 were discarded. The resulting network was composed of 79,515 nodes and 654,122 edges. The network was used to select all direct neighbors of the experimentally confirmed active compounds to have their bioactivity potential predicted. The neighbors were searched in the largest connected component of the network for the five Assays, as these would be good candidates to have their bioactivity predicted by the QSAR models. We were able to obtain 530 direct neighbors for the five Assays, from these, 102 were predicted as bioactive for one or more Assays. The similarity network is densely connected, making it hard to search for patterns visually. To enable the visualization of highly connected direct neighbors, NPMINE implements an edge filtering algorithm, leaving only the highest similarity edge by node, except when removing the edge would disconnect the network, in which case one node is allowed to have more than one edge. To illustrate this concept, one active node (compound Testosterone propionate) was selected on the network, derived from Assay AID-651838 (*qHTS assay for identifying genotoxic compounds that show differential cytotoxicity against a panel of isogenic chicken DT40 cell lines with known DNA damage response pathways*) so that the predicted bioactivity of the node’s neighbors can be inspected (Fig. 3-B).. Another significant result is the confirmation of the compound’s biological activity (other than the Bioassays [see Additional file 4] reported here) reported for 38 structures in the publications.

**Figure 3.**
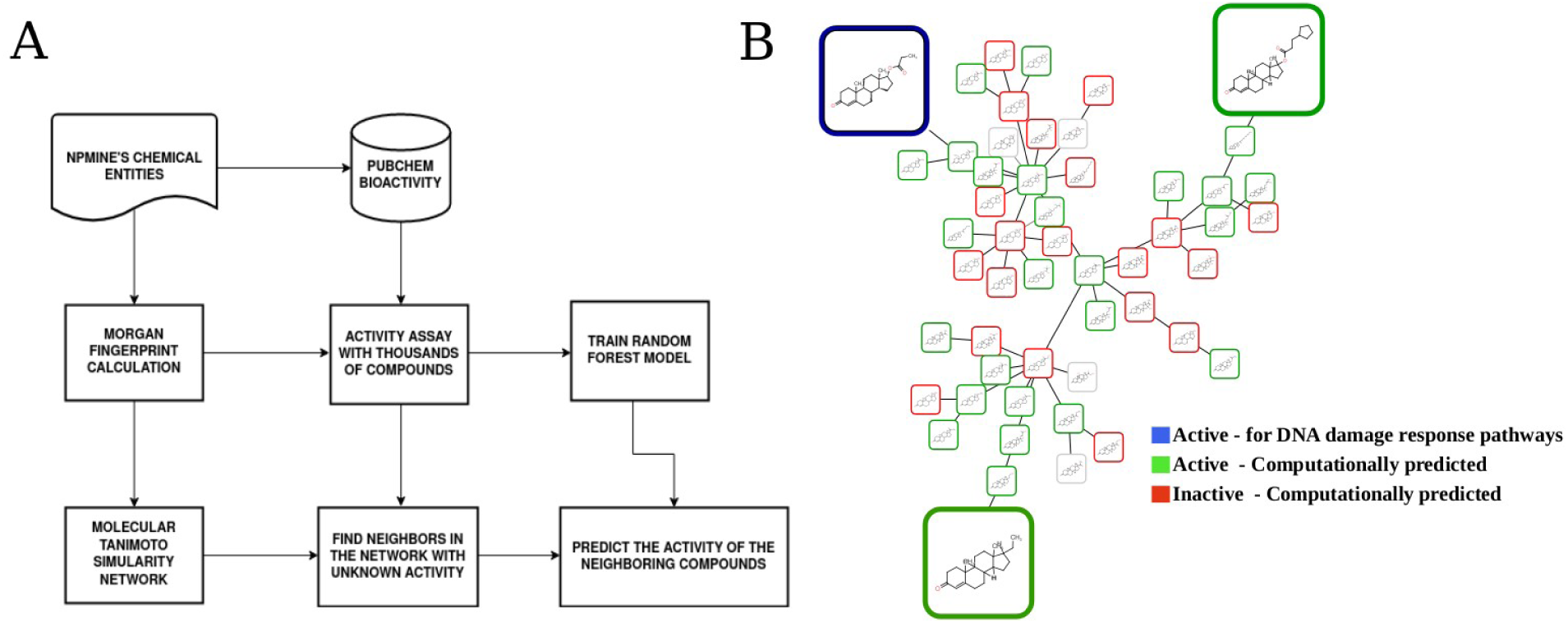
Bioactivity Prediction. A - Parallel use of QSAR and structure similarity to fit a model, find a similar structure and predict activity. B - Maximum similarity edges for each neighbor of compound Testosterone propionate (PubChem CID: 5995) highlighting structural similarity of structures used for prediction.

The functionalities to retrieve information from PubChem BioAssay database, fit QSAR models and then predict activity for new structures were incorporated into NPMINE, allowing characterization of compounds recovered from large document collections.

### Comparison with relevant natural product databases

In order to assess the representativity of NPMINE’s recovered chemical entities, they were compared with three major natural product databases, the Dictionary of Natural Products (DNP - Version 23:2A) [18], SUP [14] and the NPAtlas [15], a manually curated database (Fig. 4-A). In total 18,551 (15.34%) compounds were present in one or more natural products databases. This proportion is smaller than the number of compounds deposited in PubChem, which is 73,428 (60.7%), a factor that may be explained by the recovery of synthetic compounds, compounds with structural errors, and simply the absence in the selected databases.

**Figure 4.**
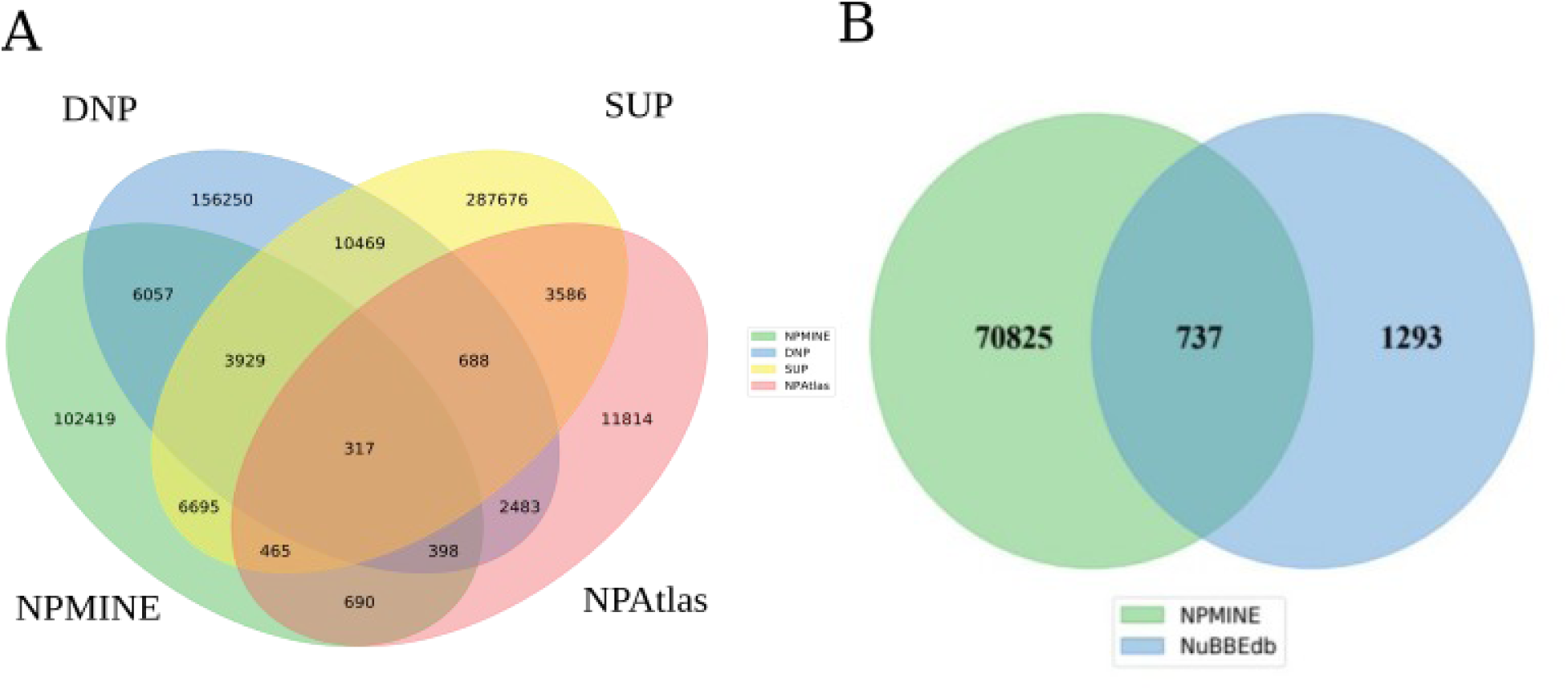
A - Venn Diagram comparing NPMINE results to natural product databases DNP, SUP and NPAtlas. B - Comparison of NPMINE results and the NUBBEDB.

The overlap with NPs databases is reassuring, as it provides evidence that the tool is indeed recovering known natural product structures automatically. The comparison with NPAtlas, a manually curated database, that assigns biological source to compounds recorded, allows the check of taxonomic assignments by NPMINE. Taking into consideration the 1870 compounds shared between NPMINE recovered and NPAtlas, counting the number of assigned genera by NPAtlas we have: 1° Streptomyces (230), 2° Aspergillus (146) and 3° Penicillium (121). Looking at the number of genera recovered (it is possible to have more than one genus by document) by Gnfinder in the same documents where these structures were recovered, we have: 1° Aspergillus (214), 2° Streptomyces (211), 3° Escherichia (208), 4° Staphylococcus (200), 5° Candida (165), 6° Bacillus (163), 7° Pseudomonas (120) e 8° Penicillium (100). The high frequency of Fungi and Bacteria associated genera indicates that NPMINE is able to provide insights on the biological source. Looking at the overlap with DNP, according to [5], the proportion of compound biological sources are 67.3% for Plantae, 12.6% for Animalia, 10% for Fungi and 9% for Bacteria. From the 10,701 shared between NPMINE recovered and DNP, when the taxonomic mentions on the same document were recovered we have the proportions Plantae 41%, Animalia 28.4%, Fungi 14,2% and Bacteria 14%, showing a correlated pattern of taxonomic association to a previously manually inspected database. Another useful comparison was performed with NuBBEDB [19]. The authors report that the database is composed of 2,223 compounds extracted from more than 1,500 articles over four years. During the initial development of NPMINE, we were able to automatically capture 737 (36.30%) of the structures from NuBBEDB (Fig. 4-B), showing the potential for speeding the cataloging process. This is evidence that, while NPMINE should not be used alone, it can be used with manual database construction as a supplementary tool.

### Taxonomic unit recovery

A large number of unique species was recovered (as shown in Table 1) and 61,6% of total unique species (accounting for overlaps between journals) were found in the Journal of Natural Products, 7,26% in Metabolomics, 39,9% in Phytochemistry Reviews, 16,4% in *Química Nova*, 13,47% and 13,8% in the Journals of Chromatography A and B, respectively. These numbers are an initial indicator that NPMINE can be used to investigate any biological system. The distribution of genera for selected kingdoms is summarized in Fig. 5-A, giving an initial summary of the range of taxa the collection of documents analyzed studies.

**Figure 5.**
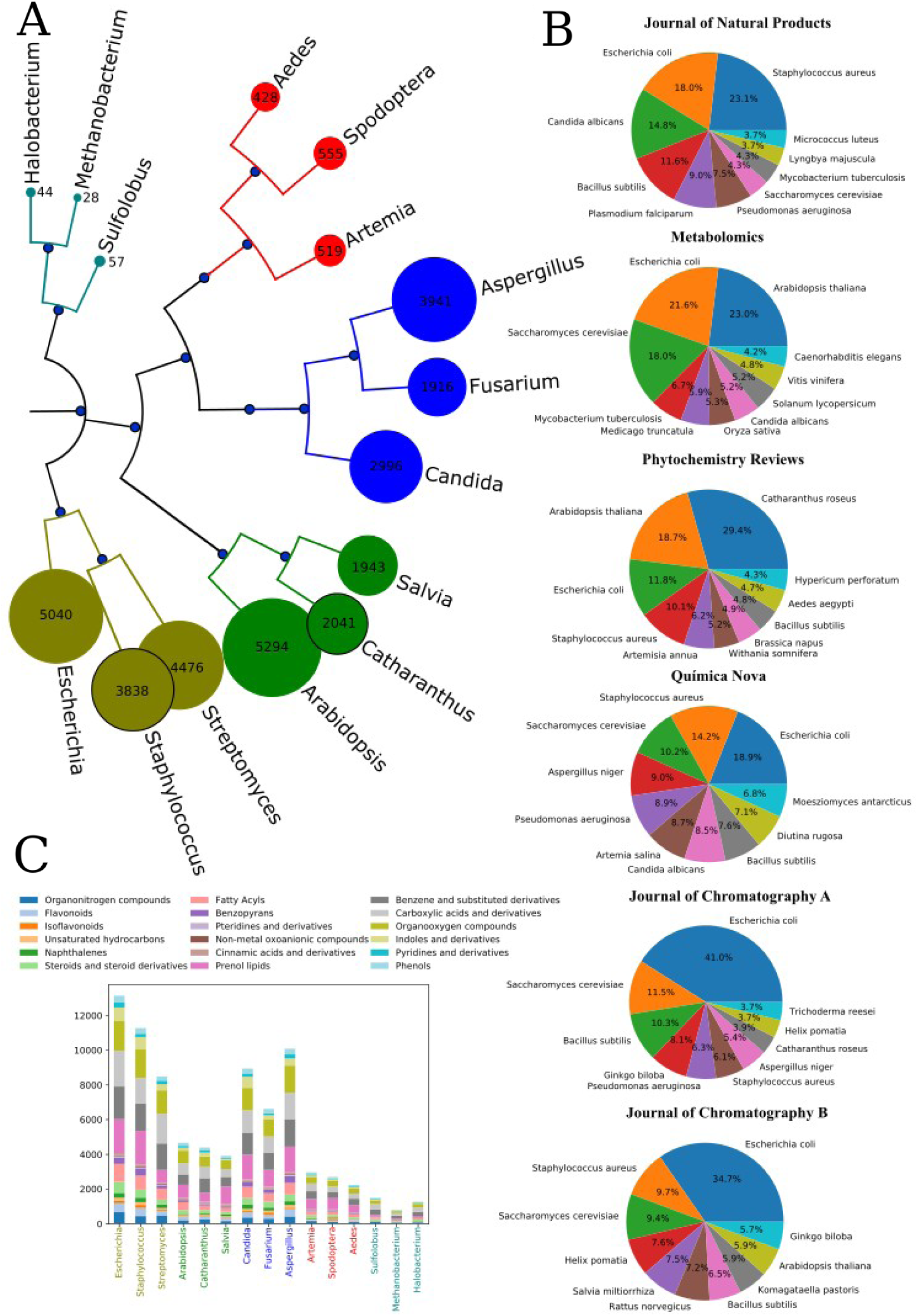
Taxonomic mentions recovered by NPMINE. A – Weighted Genera mentions recovered for selected taxonomic Kingdoms. B – Taxonomic summary by Journal. C - Number of compounds recovered in the documents where the respective genera were obtained, colored by chemical class.

For the Journal of Natural Products, the 10 most frequent species are microorganisms (Fig 5-B), mostly bacteria, which can be justified by the simplicity of experimental manipulation of microorganisms. For Metabolomics Journal (Fig. 5-B), we can see a more diverse range of species with plants, microorganisms and animals. It is clear that most species are model organisms for each kingdom, which is justified as metabolomics is a relatively new field of investigation, and basic research on the metabolism of model organisms is a reasonable starting point. One interesting aspect to be investigated on the structures found in Metabolomics, is the low novelty content, discussed above. The 10 most frequent species of Phytochemistry Reviews were not all plants, they include organisms like the mosquito *Aedes aegypti*, for which plants of different species are used as an insecticide. The plant *Catharantus roseus* is the most frequently mentioned and outranks one of the most studied plant model organisms, *Arabidopsis thaliana*. *Catharantus roseus* is a medicinal plant that possesses two invaluable antitumor terpenoid indole alkaloids called vincristine and vinblastine, but it also has international usage, as a traditional remedy for ailments such as diabetes, malaria, dengue, fever, eye irritation, diarrhea, skin infection and not only for cancer [33, 34]. In relation to the Brazilian journal *Química Nova*, it has the most even distribution of species investigated. Being a general chemistry journal, this pattern was expected. One major problem encountered in this journal was the fact that it is written in Portuguese, a Latin language, and as such, there are many commonalities between species names and Portuguese words. For example, *Figura* (figure, or graph in manuscripts) is a genus, which exemplifies the challenge to apply a tool developed to search Latin words in English manuscripts to an article in portuguese. The problem highlighted by the use of journals published in Portuguese was, in fact, widespread through the species recovered from all documents. Therefore, to include a genera from the animal kingdom we further filtered for genera including a species mentioned. For future iterations the documents have to subdivide the manuscript in sections, decreasing the possibility that author names are recovered.

Prenol lipids, that possess terpene units, are the most frequent class among compounds recovered where plant genera were mentioned (Fig. 5-C), as previously reported [5]. From the overlap between NPMINE recovered and NPAtlas, for *Streptomyces* species, the most frequent classes were Carboxylic acids and derivatives, Benzene and substituted derivatives and Organooxygen compounds.

### Study case

Using our interactive output to explore the structure/source association we focused on a narrow taxonomic group, the Orchidaceae plant family as demonstrated in the workflow found in Additional file 6 : Fig. S5. Selecting all documents containing the Orchidaceae term on the classification path variable, 1142 articles were recovered. The document IDs were used to recover the chemical entities present in these publications, recovering 3313 unique entities presenting a PubChemID. Using the PubChem BioAssay query described above, evidence of bioactivity was recorded for 1182 IDs. We finally focused on the ’Bioassay Type’ of type ‘Confirmatory’, only for compounds retrieved by OSRA. Those filters reduced our list of target compounds to 291 candidates. When these 291 candidates were inspected, we observed that many articles had mentions to Orchidaceae, without being dedicated to compounds found in this specific taxonomic group. As discussed above, these observations point to the need for more filtering steps. However, it was not difficult to recover articles dedicated to Orchidaceae bioactive related compounds. Some highlights include (±)-Homocrepidine A, Denbinobin, Gigantol, Densiflorol B and Gramistilbenoids (A, B and C), further discussed in Additional file 7: Appendix 1. NPs isolated from *Streptomyces spp.* were largely represented, while terpenoids and alkaloids were the most represented chemical classes. A large percentage of NPs (67.3%) recorded in the DNP are from the Plantae kingdom. Consistent with previous reports that approximately 70% of NP are of plant origin.

## Conclusions

In this study, we present NPMINE, a semi-automated and reproducible workflow that extracts chemical structures and taxonomic mentions from collections of PDF documents containing text and images. While previous methods such as OC processor, DECIMER, and ChemDataExtractor have also made significant contributions in extracting chemical information, NPMINE distinguishes itself by its ability to extract taxonomic names, connect and display the contextual information. Another distinct feature is the ability to mine PubChem BioAssays and predict bioactive leads on structures from literature. These features make NPMINE particularly suitable for studies related to natural products, where understanding the relationship between bioactive chemical structures and taxonomic information is crucial.

We demonstrate the efficacy of NPMINE by applying it to a collection of 55,382 scientific articles from major applied chemistry journals from Brazil and the world. Our results show consistent recovery of taxonomic and structural information from six journals, yielding 120,970 unique InChIKeys and 21,526 unique species mentions. Notably, we find that 39.3% of the identified structures were not present in PubChem, the largest public database, highlighting NPMINE’s potential for uncovering underexplored chemical structures.

Furthermore, our study reveals the bioactive potential of the entities retrieved by NPMINE. These entities exhibit predicted activity on QSAR models and exhibit substantial overlap with prominent natural product databases such as SUP, DNP, and NPAtlas. The diversity of taxonomic mentions associated with the retrieved structures suggests their applicability to a wide range of research areas, including metabolic studies and structure-bioactivity associations.

NPMINE’s versatility extends beyond scientific journals, making it suitable for academic theses and patent databases. We anticipate that our work will provide valuable resources to the scientific community, paving the way for the broader adoption and further development of NPMINE as a powerful tool for extracting chemical and taxonomic information from various sources of text and images.

## Methods

### Third-party tools

OSCAR (Open Source Chemistry Analysis Routines) is a well-established toolkit developed in 2002 for the recognition of chemical entities in scientific publications[23]. We utilized OSCAR4, the latest version of OSCAR, which offers an API for easy integration and is implemented in Java, ensuring cross-platform compatibility[23]. Given its robust functionality and portability, OSCAR4 was the tool of choice for our work.

Within NPMINE, OSCAR4 is employed as a key component of ChemExtractor, a third-party tool designed to extract chemical entities from PDF documents [36]. By leveraging the power of OSCAR4, ChemExtractor significantly accelerates the identification of chemical entities compared to manual efforts. The extracted entities are returned in a JSON (JavaScript Object Notation) format, with their corresponding InChI and InChIKeys [36].

OSRA (Optical Structure Recognition Application) is the first open-source program for optical structure recognition, benefiting from existing software components developed by the open-source software development community [11]. This collaborative approach enables continuous improvement and facilitates the program’s release as open access [11]. An important advantage of OSRA is its independence from specific document image properties, such as resolution, color depth, or font [11].

In the context of NPMINE, OSRA serves as a third-party tool to detect and extract chemical structures from scientific papers presented in graphical form. It further converts these structures into InChI and InChIKeys.

Gnfinder, also known as Global Names Finder, is a versatile third-party tool that employs Natural Language Processing (NLP) techniques to identify scientific names[24]. It operates independently and provides multiplatform packages without any external dependencies [24]. Gnfinder accepts UTF-8 encoded text as input and outputs scientific names in JSON format. Additionally, it offers the option to verify the input against various biodiversity databases through gnindex (Global Names Index) [24]. Specifically, Gnfinder utilizes NLP algorithms and complementary heuristics to detect scientific binomial or uninominal nomenclature, with our configuration focusing on English detection[37].

### Web Scraping

To gather the URL (Uniform Resource Locator) links of articles from scientific journals and therefore, apply NPMINE after downloading each one, we utilized Requests, a Python HTTP library for making requests, and BeautifulSoup, a Python HTML parsing library. Prior to employing these libraries, a comprehensive analysis of the Document Object Model (DOM) of each scientific journal website was conducted to identify the specific location of the desired information, namely the URL links of articles.

### Databases

In this work, we incorporated methods to access different databases, being these:.

1. Scientific publishers: For each specific publisher, such as the American Chemical Society, we designed scraping strategies to extract the article URL links from the standard table of contents. These links are subsequently utilized to retrieve the corresponding articles in PDF format, which serve as the basis for information retrieval. The journals included in our analysis were Journal of Natural Products, Metabolomics, Phytochemistry Reviews, Química Nova, Journal of Chromatography A, and Journal of Chromatography B.
2. Chemical structure databases: As mentioned earlier, we relied on two major databases, PubChem and ChemSpider, to associate the chemical entities extracted from scientific literature with their corresponding identifiers.
3. Biodiversity databases: these databases are accessed through Gnfinder to find the scientific names present in these articles. By default we used the NCBI and the Encyclopedia of Life databases.

### NPMINE Python package

A Python package was created to modularize each operation. Consequently, for example, if by supplying a collection of pdf documents the user wishes to only have individual access to retrieve chemical entities but not obtain the scientific names present in the specific scientific paper in question, that is possible by importing the corresponding module.

## List of Abbreviations

API: Application Programming Interface

AUC: Area Under the Curve

CACTUS: CADD Group Chemoinformatics Tools and User Services

COCONUT: Collection of Open Natural Products

DECIMER: Deep Learning for Chemical Image Recognition

DNP: Dictionary of Natural Products

DOM: Document Object Model

GNINDEX: Global Names Index

GNFINDER: Global Names Finder

HTML: HyperText Markup Language

HTTP: HyperText Transfer Protocol

INCHI: IUPAC International Chemical Identifier

IUPAC: International Union of Pure and Applied Chemistry

JSON: JavaScript Object Notation

LOTUS: Natural Products Occurance Database

NCBI: National Center for Biotechnology Information

NIST: National Institute of Standards and Technology

NLP: Natural Language Processing

NP: Natural Products

NPAtlas: Natural Products Atlas

NPMINE: Natural Product Mining

NUBBEDB: Nuclei of Bioassays, Biosynthesis and Ecophysiology of Natural Products

OSCAR: Open-Source Chemistry Analysis Routines

OSRA: Optical Structure Recognition Application

PDF: Portable Document Format

QSAR: Quantitative Structure Activity Relationship

SMILES: Simplified molecular input line entry system

URL: Uniform Resource Locator

UTF-8: 8-bit Unicode Transformation Format

SUP: Super Natural

## Availability of data and materials

All of the methods are implemented in Python and can be accessed through the GitHub repository: https://github.com/computational-chemical-biology/npmine.

## Author information

### Authors and Affiliations

**Ribeirão Preto School of Pharmaceutical Sciences, University of São Paulo, Avenida do Café, s/n, Campus da USP, Ribeirão Preto - SP, Brazil.**

Ana Carolina Lunardello Coelho & Ricardo Roberto da Silva

### Contributions

R.D.S and A.C.L.C wrote the manuscript. A.C.L.C contributed by extracting the articles and R.D.S developed the python package.

## Acknowledgments

The authors thank Camila Capel Godinho for her contribution in providing extensive knowledge and insights about Orchidaceae.

## Corresponding author

Correspondence to <u>Ricardo R. da Silva</u>.

## Supplementary Information

**Additional file 1: Fig. S1**

A representation of the generated PDF files created to benchmark NPMINE.

**Additional file 2: Fig. S2**

A comparison between the chemical structures that were submitted for benchmarking and those recovered by NPMINE, with MS being Molecular Similarity.

**Additional file 3: Table S1**

A table containing the 120,970 unique chemical compounds recovered from NPMINE applied to 55,382 articles and considering only InChI Keys.

**Additional file 4: Fig. S3**

Information on the five assays from the PubChem BioAssay database used to train QSAR models.

**Additional file 5: Fig. S4**

A representation of a chemical compound that was extracted by NPMINE in PubChem Sketcher.

**Additional file 6: Fig. S5**

A workflow demonstrating the use of NPMINE to explore structure/source associations using the plant family Orchidaceae as an example. From the 22,394 articles mentioning Orchidaceae, Denbinoben was selected as one of the compounds of interest amongst others such as Densifloral B.

**Additional file 7: Appendix 1**

Information about the Orchidaceae bioactive related compounds ((±)-Homocrepidine A, Denbinobin, Gigantol, Densiflorol B and Gramistilbenoids (A, B and C)) resulted from the workflow shown in Additional file 6.

## References

1. Newman DJ, Cragg GM (2016) Natural Products as Sources of New Drugs from 1981 to 2014. J Nat Prod 79:629–661. https://doi.org/10.1021/acs.jnatprod.5b01055

2. Harvey AL, Edrada-Ebel R, Quinn RJ (2015) The re-emergence of natural products for drug discovery in the genomics era. Nat Rev Drug Discov 14:111–129. https://doi.org/10.1038/nrd4510

3. Sparks TC, Wessels FJ, Lorsbach BA, et al (2019) The new age of insecticide discovery-the crop protection industry and the impact of natural products. Pestic Biochem Physiol 161:12– 22. https://doi.org/10.1016/j.pestbp.2019.09.002

4. Mahesh SK, Fathima J, Veena VG (2019) Cosmetic Potential of Natural Products: Industrial Applications. In: Swamy MK, Akhtar MS (eds) Natural Bio-active Compounds: Volume 2: Chemistry, Pharmacology and Health Care Practices. Springer, Singapore, pp 215–250

5. Chassagne F, Cabanac G, Hubert G, et al (2019) The landscape of natural product diversity and their pharmacological relevance from a focus on the Dictionary of Natural Products®. Phytochem Rev 18:601–622. https://doi.org/10.1007/s11101-019-09606-2

6. Pye CR, Bertin MJ, Lokey RS, et al (2017) Retrospective analysis of natural products provides insights for future discovery trends. Proc Natl Acad Sci 114:5601–5606. https://doi.org/10.1073/pnas.1614680114

7. Swain MC, Cole JM (2016) ChemDataExtractor: A Toolkit for Automated Extraction of Chemical Information from the Scientific Literature. J Chem Inf Model 56:1894–1904. https://doi.org/10.1021/acs.jcim.6b00207

8. Akhondi SA, Muresan S, Williams AJ, Kors JA (2015) Ambiguity of non-systematic chemical identifiers within and between small-molecule databases. J Cheminformatics 7:54. https://doi.org/10.1186/s13321-015-0102-6

9. Rajan K, Brinkhaus HO, Agea MI, et al (2023) DECIMER.ai - An open platform for automated optical chemical structure identification, segmentation and recognition in scientific publications

10. Barnabas SJ, Böhme T, Boyer SK, et al (2022) Extraction of chemical structures from literature and patent documents using open access chemistry toolkits: a case study with PFAS. Digit Discov 1:490–501. https://doi.org/10.1039/D2DD00019A

11. Filippov IV, Nicklaus MC (2009) Optical Structure Recognition Software To Recover Chemical Information: OSRA, An Open Source Solution. J Chem Inf Model 49:740–743. https://doi.org/10.1021/ci800067r

12. Sorokina M, Steinbeck C (2020) Review on natural products databases: where to find data in 2020. J Cheminformatics 12:20. https://doi.org/10.1186/s13321-020-00424-9

13. Sorokina M, Merseburger P, Rajan K, et al (2021) COCONUT online: Collection of Open Natural Products database. J Cheminformatics 13:2. https://doi.org/10.1186/s13321-020-00478-9

14. Banerjee P, Erehman J, Gohlke B-O, et al (2015) Super Natural II--a database of natural products. Nucleic Acids Res 43:D935–939. https://doi.org/10.1093/nar/gku886

15. van Santen JA, Jacob G, Singh AL, et al (2019) The Natural Products Atlas: An Open Access Knowledge Base for Microbial Natural Products Discovery. ACS Cent Sci 5:1824– 1833. https://doi.org/10.1021/acscentsci.9b00806

16. Lyu C, Chen T, Qiang B, et al (2021) CMNPD: a comprehensive marine natural products database towards facilitating drug discovery from the ocean. Nucleic Acids Res 49:D509– D515. https://doi.org/10.1093/nar/gkaa763

17. Afendi FM, Okada T, Yamazaki M, et al (2012) KNApSAcK Family Databases: Integrated Metabolite–Plant Species Databases for Multifaceted Plant Research. Plant Cell Physiol 53:e1. https://doi.org/10.1093/pcp/pcr165

18. Dictionary of Natural Products 30.2 Chemical Search. https://dnp.chemnetbase.com/faces/chemical/ChemicalSearch.xhtml. Accessed 22 Mar 2022

19. Pilon AC, Valli M, Dametto AC, et al (2017) NuBBEDB: an updated database to uncover chemical and biological information from Brazilian biodiversity. Sci Rep 7:7215. https://doi.org/10.1038/s41598-017-07451-x

20. Ntie-Kang F, Zofou D, Babiaka SB, et al (2013) AfroDb: A Select Highly Potent and Diverse Natural Product Library from African Medicinal Plants. PLOS ONE 8:e78085. https://doi.org/10.1371/journal.pone.0078085

21. Rutz A, Sorokina M, Galgonek J, et al (2021) The LOTUS Initiative for Open Natural Products Research: Knowledge Management through Wikidata. 2021.02.28.433265

22. Köster J, Rahmann S (2012) Snakemake—a scalable bioinformatics workflow engine. Bioinformatics 28:2520–2522. https://doi.org/10.1093/bioinformatics/bts480

23. Jessop DM, Adams SE, Willighagen EL, et al (2011) OSCAR4: a flexible architecture for chemical text-mining. J Cheminformatics 3:41. https://doi.org/10.1186/1758-2946-3-41

24. Mozzherin D, Myltsev A, Zalavadiya H (2022) gnames/gnfinder: v0.18.2. Zenodo

25. LaTeX - A document preparation system. https://www.latex-project.org/. Accessed 22 Mar 2022

26. Federhen S (2012) The NCBI Taxonomy database. Nucleic Acids Res 40:D136–D143. https://doi.org/10.1093/nar/gkr1178

27. Rajan K, Brinkhaus HO, Zielesny A, Steinbeck C (2020) A review of optical chemical structure recognition tools. J Cheminformatics 12:60. https://doi.org/10.1186/s13321-020-00465-0

28. Ihlenfeldt WD, Bolton EE, Bryant SH (2009) The PubChem chemical structure sketcher. J Cheminformatics 1:20. https://doi.org/10.1186/1758-2946-1-20

29. Yonchev D, Dimova D, Stumpfe D, et al (2018) Redundancy in two major compound databases. Drug Discov Today 23:1183–1186. https://doi.org/10.1016/j.drudis.2018.03.005

30. Vazquez M, Krallinger M, Leitner F, Valencia A (2011) Text Mining for Drugs and Chemical Compounds: Methods, Tools and Applications. Mol Inform 30:506–519. https://doi.org/10.1002/minf.201100005

31. NCI/CADD Group Chemoinformatics Tools and User Services. https://cactus.nci.nih.gov/. Accessed 22 Mar 2022

32. Pedregosa F, Varoquaux G, Gramfort A, et al (2011) Scikit-learn: Machine Learning in Python. J Mach Learn Res 12:2825–2830

33. Nejat N, Valdiani A, Cahill D, et al (2015) Ornamental exterior versus therapeutic interior of Madagascar periwinkle (Catharanthus roseus): the two faces of a versatile herb. ScientificWorldJournal 2015:982412. https://doi.org/10.1155/2015/982412

34. Goboza M, Aboua YG, Chegou N, Oguntibeju OO (2019) Vindoline effectively ameliorated diabetes-induced hepatotoxicity by docking oxidative stress, inflammation and hypertriglyceridemia in type 2 diabetes-induced male Wistar rats. Biomed Pharmacother 112:108638. https://doi.org/10.1016/j.biopha.2019.108638

35. Barnabas S, Böhme T, Boyer S, et al (2022) Extraction of Chemical Structures from Literature and Patent Documents using Open Access Chemistry Toolkits: A Case Study with PFAS. https://doi.org/10.26434/chemrxiv-2022-nmnnd-v3

36. Williamson MJ (2022) Chemextractor

37. Rutz A, Dounoue-Kubo M, Ollivier S, et al (2019) Taxonomically Informed Scoring Enhances Confidence in Natural Products Annotation. Front Plant Sci 10:

38. Heller S, McNaught A, Stein S, et al (2013) InChI - the worldwide chemical structure identifier standard. J Cheminformatics 5:7. https://doi.org/10.1186/1758-2946-5-7

